# Complementary, Semi-automated Methods for Creating Multi-dimensional, PEG-based Biomaterials

**DOI:** 10.1101/219501

**Authors:** Elizabeth A. Brooks, Lauren E. Jansen, Maria F. Gencoglu, Annali M. Yurkevicz, Shelly R. Peyton

**Affiliations:** Department of Chemical Engineering, University of Massachusetts Amherst, N540 Life Sciences Laboratories, 240 Thatcher Rd., Amherst, MA 01003-9364.

**Keywords:** hydrogels, liquid handling robotics, extracellular matrix, breast cancer

## Abstract

Tunable biomaterials that mimic selected features of the extracellular matrix (ECM), such as its stiffness, protein composition, and dimensionality, are increasingly popular for studying how cells sense and respond to ECM cues. In the field, there exists a significant trade-off for how complex and how well these biomaterials represent the *in vivo* microenvironment, versus how easy they are to make and how adaptable they are to automated fabrication techniques. To address this need to integrate more complex biomaterials design with high-throughput screening approaches, we present several methods to fabricate synthetic biomaterials in 96-well plates and demonstrate that they can be adapted to semi-automated liquid handling robotics. These platforms include 1) glass bottom plates with covalently attached ECM proteins, and 2) hydrogels with tunable stiffness and protein composition with either cells seeded on the surface, or 3) laden within the three-dimensional hydrogel matrix. This study includes proof-of-concept results demonstrating control over breast cancer cell line phenotypes via these ECM cues in a semi-automated fashion. We foresee the use of these methods as a mechanism to bridge the gap between high-throughput cell-matrix screening and engineered ECM-mimicking biomaterials.

## INTRODUCTION

Synthetic biomaterials are valuable platforms for studying cell behavior *in vitro* because they capture some features of real tissue microenvironments. These factors are missed using standard tissue culture polystyrene (TCPS) and are not easily modified using *in vivo* models. Many have used engineered biomaterials to better understand biological processes such as organoid development,^1^ stem cell differentiation,^2-3^ tumor progression,^4-5^ and drug response.^6-7^ Bioengineers in the field have developed many different classes of biomaterial systems in which to study cell-extracellular matrix (ECM) interactions. These can be protein or peptide-based systems, or made from synthetic polymer precursors.^8^ We focus here on this latter class of materials because they are engineered from the ground-up, and provide a plethora of tunable characteristics to direct cell function. These biomaterials have been developed to either allow for cell culture on their surfaces (Two-dimensional (2D) hydrogels), or to allow for encapsulation within the material (Three-dimensional (3D) hydrogels). 3D hydrogel models have been used to independently tune biophysical and biochemical cues to determine their impact on intestinal stem cell organoid growth,^1^ cancer cell invasion,^9^ and drug response.^10^ Moreover, the dynamics of microenvironments have been captured in some 3D models by using photoinitiated polymerization to change the hydrogel modulus and ligand density^11^ and by creating patterns within a hydrogel to control the spatial arrangement of biochemical cues within the matrix to direct cell phenotype.^12^ Lastly, more complex material designs that include microfluidics have been used to capture the bone marrow,^13^ liver,^14^ and heterogeneous tumor microenvironments,^15-16^ or to study the response of lung inflammation^17^ and glioblastoma^18^ to drugs. While these systems capture fluid flow and spatial gradients, they are generally labor-intensive^18^ and low-throughput,^17^ which makes expansion to large-scale studies a significant challenge.

One area of interest is using biomaterials to make preclinical and clinical predictions of diseases. For example, our group demonstrated that breast cancer cell speed, displacement, and adhesivity were dependent on the ECM protein composition coupled to glass, and these phenotypes could be used to predict tissue-specific breast cancer metastasis. We and others have demonstrated drug that response is affected by both stiffness and ECM protein composition on 2D hydrogels.^6-7, 20^ We have also shown that tumor spheroids encapsulated into 3D hydrogels are drug resistant compared to TCPS.^10^ Additionally, biomaterials can be used to predict stage-specific phenotypes in disease. For example, early stage melanoma cell drug response was shown to be insensitive to the dimensionality of the platform, but metastatic melanoma cells were resistant in 3D environments.^21^ These studies demonstrate controlled *in vitro* ECMs are useful tools for preclinical studies.

The need for engineered biomaterials in the drug development pipeline is clear, since only 8% of filed Investigational New Drug applications will be approved by the Food and Drug Administration.^22^ This is due in part to false positives missed early in the drug development pipeline. Therefore, there is a critical need to scale up these types of biomaterials in order for them to be feasible for high-throughput screening. To achieve a paradigm shift in high-throughput screening with biomaterials, they must be formatted to use existing equipment and technology, such as liquid handling robotics and plate readers.^23^

To address this need, we integrated three biomaterials our lab has developed into 96-well plates and used semi-automated liquid handling robotics to fabricate these platforms (Figure 1). In this study, we give a detailed overview of their development, use, and how they can be made amenable to semiautomated technologies. They range from the simplest, chemically modified glass surfaces, to a significantly more complex tunable 3D hydrogel cell culture system. We propose that each of these platforms are incredibly useful for *in vitro* studies of cell-ECM interactions, and the level of complexity chosen by an individual lab should be guided by the biological question at hand.

**Figure 1.**
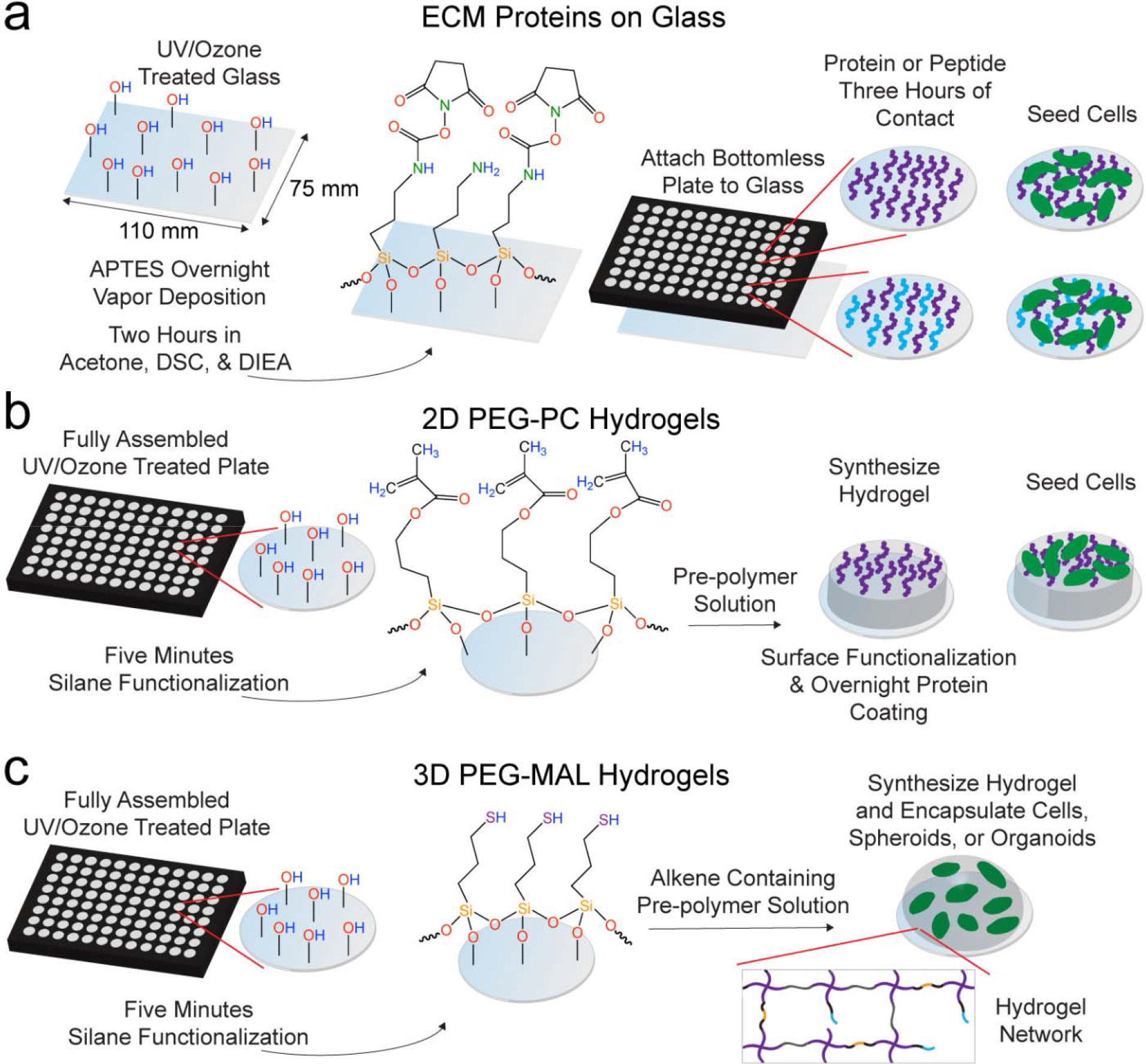
Schematic of three high-throughput, 96-well biomaterial platforms for studying cell behavior *in vitro*. a. To make protein-modified glass-bottom plates, we chemically modified a well plate-sized piece of glass. This glass was attached to a bottomless 96-well plate and ECM proteins (purple and blue) were attached in solution with three hours of surface contact followed by a PEG blocking step for two hours before cell (green) seeding. b. 2D PEG-PC hydrogels were covalently attached to fully assembled glass bottom 96-well plates functionalized with a methacrylate silane. The hydrogels were functionalized with full length proteins via sulfo-SANPAH before cell seeding. c. 3D PEG-MAL hydrogels were covalently attached to fully assembled glass bottom 96-well plates functionalized with MPT. The PEG-MAL (4-arm, purple) pre-polymer solution containing cell-adhesion (black/blue) and cell-degradable (black/orange) peptides was mixed with single cells and the PEG-dithiol crosslinker (gray), and allowed to polymerize on the silane-functionalized glass surface.

## MATERIALS AND METHODS

### Cell culture

All cell culture supplies were purchased from Thermo Fisher Scientific (Waltham, MA). MDA-MB-231 and Hs578T breast cancer cell lines were grown in Dulbecco’s Modified Eagle Medium (DMEM), supplemented with 10% fetal bovine serum (FBS) and 1% penicillin/streptomycin (pen/strep) at 37°C and 5% CO_2_. The MDA-MB-231 cell line was provided by Dr. Sallie Smith Schneider at the Pioneer Valley Life Sciences Institute, and Hs578T cells were provided by Dr. Mario Niepel at Harvard Medical School.

### Custom 96-well plate fabrication

We used bottomless polystyrene 96-well plates (Greiner Bio-One, Kremsmünster, Austria) as the base of the custom well plate. We manually cut a rectangular piece of ARcare 90106 (Adhesives Research, Glen Rock, PA)^24^ medical-grade, double-sided adhesive to fit the well plate bottom, and it was attached to the plate by hand ensuring that there were no air bubbles present. Holes were cut in the adhesive from a custom template designed in Adobe Illustrator CC 2014 (Adobe Systems, San Jose, CA) with a laser cutter (Epilog Mini 18 Laser, Golden, CO). We purchased glass pieces cut to size (110 mm x 75 mm, S.I. Howard Glass, Worcester, MA) with the same thickness as a number 1.5 coverslip (0.17 mm) to optimize microscopy applications. We adhered the glass to the plate by removing the remaining backing to the adhesive and pressing the glass firmly to ensure a tight seal around each well. The glass was attached to the plate before or after chemical modification, depending on the biomaterial platform (Figure S1).

### Basic liquid handling robotics operation

We have a Biomek NX^P^ Liquid Handling Automation Workstation (Biomek, Beckman Coulter, Brea, CA) configured with 250 μL syringe pumps. This instrument is enclosed in a 1.0 m x 0.9 m Biomek NX^P^ Negative Pressure HEPA Enclosure (Beckman Coulter) to enable sterile cell culture. With the instrument powered on, we ran the Biomek software (version 4.1) on the computer connected to the Biomek to control the instrument. Each day of an experiment, we ran the “Home All Axes” method until we observed that no air bubbles were present in the syringes. Individual methods were created for each platform and experiment with the Biomek software. We primarily used the “Aspirate” and “Dispense” functions within the software to perform protein solution dilutions and to pipette the 2D and 3D hydrogel pre-polymer solutions into our 96-well plate platforms.

### ECM protein well plates

ECM proteins were attached to the glass surface in a 96-well plate format by modifying a procedure described previously.^19^ For each plate, one 110 mm x 75 mm piece of glass was UV/ozone treated at 1 atmosphere (UV/Ozone ProCleaner, 180 micrograms/m^3^ ozone level in chamber, 4.6 mW/cm^2^ peak UV intensity, 10.15 W UV lamp power requirement, BioForce Nanosciences, Salt Lake City, UT) for 10 minutes to hydroxylate the surface. The glass was placed in an inverted well plate lid supported by small acrylic posts to allow for air flow on both sides of the glass. (3-aminopropyl)triethoxysilane (APTES, Sigma-Aldrich, St. Louis, MO) was deposited into inverted polypropylene lids spaced evenly around the glass. Another well plate lid was placed on top and this container was sealed with autoclave tape and wrapped in aluminum foil. This was placed in a 90°C oven to functionalize the glass with APTES by vapor deposition overnight as previously described. The glass was rinsed four times sequentially in toluene (Thermo), 95% ethanol (Pharmco-AAPER, Brookfield, CT), and water. The glass was dried at 90°C for one hour and then functionalized with 10 g/L N,N-disuccinimidyl carbonate (DSC, Sigma-Aldrich) and 5 v% N,N-Diisopropylethylamine (DIEA, Sigma-Aldrich) in acetone (Thermo) for two hours. The glass was rinsed three times in acetone, air-dried, and used immediately or stored in a desiccator overnight. The surface of each well was coated with a 40 µL protein solution containing 0, 0.13, 0.25, 0.5, 1, or 2 μg/cm^2^ collagen I (rat tail, Thermo) with or without the addition of a constant 0.5 μg/cm^2^ fibronectin (human, Millipore, Billerica, MA) background diluted in pH 7.4 phosphate buffered saline (PBS) by manual pipetting or with liquid handling robotics (Biomek). The plates were rinsed three times with PBS and the remaining surface area was blocked with 10 μg/cm^2^ MA(PEG)_24_ (Thermo) for two hours to prevent non-specific protein adsorption. The wells were rinsed three times, and UV sterilized for one hour before cell seeding. MDA-MB-231 or Hs578T cells were seeded in serum free cell culture medium at 3,400 cells/cm^2^ and were imaged with an Axio Observer Z1 (Carl Zeiss AG, Oberkochen, Germany) one day post-seeding, N ≥ 85 cells were analyzed per condition.

### X-ray photoelectron spectroscopy (XPS) analysis

Glass coverslips (15 mm diameter, number 1.5, Thermo) were treated with UV/ozone, APTES, DSC, and 2 μg/cm^2^ of the cell-adhesion peptide GSPCRGDG (RGD, Genscript, Piscataway, NJ) as described above. Coverslips were used instead of large pieces of glass to fit into the instrument for analysis. Coverslips representative of each treatment step were analyzed by XPS (Physical Electronics Quantum 2000 Scanning ESCA microprobe, Physical Electronics, Inc., Chanhassen, MN). Spectra were obtained with a 200 µm spot size and monochromatic Al Kα radiation (1,486.68 eV). The x-ray source was operated at 45.9 W with the analyzer’s constant pass energy at 187.85 eV. The take off angle was 45° from the sample surface and N = 2 locations per sample.

### 2D hydrogel platform

2D poly(ethylene glycol)-phosphorylcholine (PEG-PC) hydrogels were synthesized using modifications to a method described previously.^6^ Briefly, fully assembled glass bottom 96-well plates were UV/ozone treated (BioForce Nanosciences) for 10 minutes. A 2 v% solution of 3-(trimethoxysilyl)propyl methacrylate (MPS, Sigma-Aldrich) in 95% ethanol (adjusted to pH 5.0 with glacial acetic acid) was used to silane-functionalize the wells for five minutes. The wells were rinsed three times with 100% ethanol and dried in an oven. A 17 wt% solution of 2-methacryloyloxyethyl phosphorylcholine (PC, Sigma-Aldrich) in pH 7.4 PBS was mixed with 1, 3, 6, and 10 wt% poly(ethylene glycol) dimethacrylate (PEGDMA, Mn 750, Sigma-Aldrich). These PEG-PC solutions were clarified by filtering through a 0.2 µm syringe filter (Thermo). Hydrogel polymerization was initiated by lithium phenyl(2,4,6-trimethylbenzoyl)phosphinate (LAP, TCI America, Portland, OR).^26^ LAP at 200 mM dissolved in 70% ethanol was added to the pre-polymer solutions to a final concentration of 2 mM to polymerize the hydrogels.^26^ PEG-PC hydrogels were formed either manually using a multi-channel pipette or with liquid handling robotics (Biomek) by adding 40 μL of the pre-polymer solution to the wells and placing the plates under UV light (365 nm, UV Panel HP, American DJ, Los Angeles, CA) to induce hydrogel polymerization for at least 20 minutes. Post-polymerization, 100 μL PBS was added to each well to allow the hydrogels to swell overnight. Fully swollen hydrogels ranged from 500 to1000 μm thick depending on the ratio of PEG to PC.

The swollen hydrogels were functionalized with protein by reacting with 0.6 mg/mL sulfosuccinimidyl 6-(4′-azido-2′-nitrophenylamino)hexanoate (sulfo-SANPAH, ProteoChem, Denver, CO) dissolved in (4-(2-hydroxyethyl)-1-piperazineethanesulfonic acid) (HEPES) buffer (pH 8.5) under 365 nm UV light for at least 20 minutes. The hydrogels were rinsed three times with HEPES buffer. Protein solutions in pH 3.8 PBS were added to the wells for a final protein concentration of 5 μg/cm^2^ containing either collagen I alone (Thermo) or a “collagen-rich” solution containing 65% collagen I (Thermo), 33% collagen III (recombinant human, FibroGen, San Francisco, CA), and 2% fibronectin (Millipore).^6^ The plates were stored at 4°C overnight and rinsed three times with pH 3.8 PBS, followed by three rinses with pH 7.4 PBS, UV sterilized for one hour, and then rinsed with cell culture medium prior to cell seeding. MDA-MB-231 cells were seeded at 34,000 cells/cm^2^ and Hs578T cells were seeded at 17,000 cells/cm^2^.

### 3D hydrogel platform

Glass bottom 96-well plates were UV/ozone treated (BioForce Nanosciences) for 10 minutes and functionalized with 2 v% solution of (3-mercaptopropyl)trimethoxysilane (MPT, Thermo) in 95% ethanol (adjusted to pH 5.0 with glacial acetic acid) for five minutes. The wells were rinsed three times with 100% ethanol and dried in an oven. For mechanical testing and hydrogel survival studies no cells were encapsulated into the hydrogels. A 4-arm PEG-maleimide (PEG-MAL, 20 kDa 4-arm PEG-MAL, Jenkem Technology, Plano, TX) was dissolved at 5, 10, or 20 wt% in serum free DMEM and crosslinked at a 1:1 ratio with linear PEG-dithiol (PDT, 1 kDa, Sigma-Aldrich) dissolved in 2 mM triethanolamine (pH 7.4) as a catalyst.^27^ Final hydrogel volume was 10 µL, which covers the surface of the well once swollen. After waiting five minutes to ensure hydrogels were completely polymerized, the hydrogels were swollen overnight by adding 100 μL/well PBS. Fully swollen hydrogels were approximately 500 μm thick. Cells were encapsulated into 3D PEG-MAL hydrogels as described previously^10^ with minor modifications. Briefly, a 10 wt% solution of PEG-MAL containing 2 mM of cell-adhesion peptide RGD (GRGDSPCG, Genscript), GFOGER (CGP(GPP)_5_GFOGER(GPP)_5_, Genscript), DGEA (GCGDGEA, see the Supplemental Information), or no cell-adhesion peptide was combined with either MDA-MB-231 or Hs578T cells at 500,000 cells/mL. This solution was crosslinked at a 1:1 with linear PDT (1 kDa, Sigma-Aldrich) or a 1:1 molar combination of the linear PDT and cell-degradable crosslinker, Pan-matrix metalloproteinase (Pan-MMP, GCRDGPQGIWGQDRCG, Genscript) in sterile 2 mM triethanolamine. The hydrogels were synthesized by pipetting 1 μL of the PDT or PDT and Pan-MMP solution onto the well surface manually or with the liquid handling robotics followed by pipetting 9 μL of the cell-containing PEG-MAL solution manually near the well surface or with the liquid handling robotics at 5 mm above the well surface to prevent polymerization of the hydrogels at the pipette tips.

### Collagen I ELISA

Plates were prepared as described above with solutions containing 0-2 µg/cm collagen I alone or with an additional 0.5 μg/cm fibronectin background with manual pipetting. The plates were blocked with non-specific protein by incubating overnight at 4°C with AbDil (2 wt% bovine serum albumin (Thermo) in tris-buffered saline (TBS) with 0.1% Triton x-100 (Sigma-Aldrich), TBS-T). The AbDil was removed and a 1:200 dilution of a collagen I antibody (ab6308, Abcam, Cambridge, MA) in AbDil was added to each well and incubated for one hour while rotating the plate. The primary antibody was removed and the wells were rinsed three times with washing buffer (0.5% Tween 20 (Sigma-Aldrich) in PBS). The secondary anti-mouse HRP antibody (PI31430, Thermo) was added at a 1:20,000 dilution in AbDil for one hour while rotating the aluminum foil-wrapped plate. The antibody was removed and the wells were rinsed three times with washing buffer and then 50 μL of Pierce 1-Step Ultra TMB-ELISA Substrate Solution (PI34028, Thermo) was added to each well. The plate was incubated in the dark for 15 minutes. The reaction was quenched by adding 50 μL/well 2 M sulfuric acid. The absorbance was read at 450 nm (BioTek ELx800 microplate reader, BioTek, Winooski, VT). For the plotted data for the collagen I only ELISA, N = 2 independent experiments and n = 3 technical replicates. For the collagen I with fibronectin background, n = 3 technical replicates. The data are plotted as the mean ± standard error of the mean (SEM) in both cases.

### Hydrogel mechanical characterization

A custom-built instrument described previously^28^ was used to measure the effective Young’s modulus of the 2D PEG-PC and 3D PEG-MAL hydrogels made in 96-well plates (manual or semi-automated pipetting) and on coverslips (manual pipetting). Here, indentation testing was performed on 40 μL 2D hydrogels and 10 μL 3D hydrogels. In brief, a flat punch cylindrical probe (custom made using a High-Speed M2 Tool Steel Hardened Undersized Rod from McMaster Carr, Elmhurst, IL, diameter 3.1-5.2 mm) was brought into contact with the surface of the hydrogel at a rate of 10 μm/s to a maximum displacement of 100 μm. The sample height (h) to the contact radius (a) was kept between 0.5<a/h<2 and material compliance was analyzed using a correction ratio between the contact radius and the sample height to account for the dimensional confinement, as previously described.^29^ This modified Hertzian model was used to analyze the first 10% of the linear portion of the force-indentation curve.^29^ For mechanical characterization, N ≥ 4 hydrogels per condition.

### Hydrogel survival and silane stability curves

For the hydrogel survival curves, 2D PEG-PC and 3D PEG-MAL hydrogels were synthesized as described above in 96-well plates (manual or semiautomated pipetting) or on coverslips (manual pipetting) and swollen in PBS. The PBS was replenished/changed two to three times per week and hydrogel covalent attachment was verified with plate inversion and visual inspection. For the hydrogel survival curves, N ≥ 6 hydrogels per condition. For the silane chemistry stability, 15 mm diameter glass coverslips were prepared with the MPT chemistry described above and stored in a 24-well plate on the bench or in a desiccator. At each time point, three new 10 wt% PEG-MAL hydrogels were synthesized manually with 1:1 crosslinking with linear PDT. PBS was added to each well and hydrogel adherence was determined by visual inspection after five minutes.

### Immunofluorescence staining

Breast cancer cells seeded on TCPS or 2D PEG-PC hydrogels (10 wt% PEGDMA) were fixed and stained one day post-seeding. Cells were rinsed three times with warm PBS, fixed with 10% formalin (Thermo) at 37°C for 20 minutes. Cells were rinsed again three times with PBS, permeabilized with TBS + 0.5% Triton x-100 for 10 minutes, rinsed three times with PBS and blocked in AbDil for 20 minutes. A 1:300 dilution of a monoclonal mouse anti-vinculin antibody (V9264, Sigma-Aldrich) in AbDil was added to the wells for one hour while the plate was gently rotated, removed, and the cells were rinsed three times with TBS-T. A 1:500 dilution of Alexa Fluor anti-mouse 555 (Thermo) was added for 45 minutes while the aluminum foil-wrapped plate was gently rotated. The hydrogels were rinsed three times with TBS-T. To stain for F-actin, 1:40 dilution of Alexa Fluor 647 phalloidin (Thermo) in PBS was added to the wells for 20 minutes while gently rotating the aluminum foil-wrapped plate. The cells were rinsed three times with PBS. The nuclei were stained with a 1:10,000 dilution of DAPI (Thermo) in PBS for five minutes. The cells were rinsed three times with PBS. Cells were imaged using a Zeiss Spinning Disc Cell Observer SD (Zeiss). The images shown are representative from a set of N = 2 independent experiments.

### LIVE/DEAD cell viability staining

At one and four days after encapsulation in 3D PEG-MAL hydrogels, cells were evaluated for viability by LIVE/DEAD staining (L3224, Thermo) in accordance with the manufacturer’s instructions. Cells in the hydrogels were imaged using a Zeiss Spinning Disc Cell Observer SD (Zeiss). The results were quantified by manually counting the cells in the images, N ≥ 6.

#### Image processing

Cell morphology was evaluated by manual tracing in ImageJ (National Institutes of Health, Bethesda, MD). For cells on ECM well-plates, N ≥ 85. The LIVE/DEAD image stacks of the cells encapsulated in the 3D hydrogels were converted to 2D images in Imaris ×64 7.7.2 (Bitplane, Belfast, United Kingdom) using the Easy 3D display mode. The results were quantified by manually counting the cells in the images, N ≥ 6.

#### Statistical analysis

Statistical analysis was performed with GraphPad Prism v7.0c (GraphPad Software, Inc., LaJolla, CA). Statistical significance was evaluated using a one-way analysis of variance (ANOVA) followed by a Tukey’s post-test for pairwise comparisons. For the analysis, p-values <0.05 are considered significant, where p<0.05 is denoted with *, ≤0.01 with **, ≤0.001 with ***, and ≤0.0001 with ****

## RESULTS

### Three distinct biomaterial platforms adapted to 96-well plates and liquid handling robotics

We previously developed a biomaterial system with ECM proteins covalently attached to glass, providing control over the integrin-binding of the cells by changing the protein composition on the surface.^19^ This chemistry was initially developed for single coverslips, and this method was incompatible with semiautomated applications. To optimize this chemistry for multi-well plates, we applied this same silane chemistry on a well plate-sized piece of glass (Figure 1a). We used UV/ozone treatment to promote surface hydroxylation followed by overnight chemical vapor deposition of APTES. We functionalized the surface with DSC to provide the amine-reactive succinimidyl ester and then attached the glass to a bottomless 96-well plate with a medical grade adhesive (Figure S1a-b). Then, the desired single protein or protein combination was covalently attached within each well using semi-automated liquid handling robotics. The remaining surface area was blocked with non-specific PEG to ensure that the cells would only bind to the intended proteins on the surface. Then, cells were seeded into the wells for experiments.

2D PEG-PC hydrogels were synthesized in 96-well plates with an efficient photoinitiator (Figure 1b). We improved this platform from our initial adaptation in a 96-well plate^6^ by using semi-automated liquid handling robotics to pipette the pre-polymer solution of the 2D PEG-PC hydrogels, and by initiating the polymerization with UV light in a 96-well plate. Fully assembled glass bottom 96-well plates were UV/ozone treated and then functionalized with a solution of MPS. This silane functionalization on the glass surface reacted with the methacrylates in the PEGDMA, which covalently attached the 2D hydrogels to the glass during polymerization. We tuned the hydrogel stiffness by varying the mesh size of the hydrogel network, and these hydrogels were functionalized with proteins via the amine-reactive sulfo-SANPAH before seeding cells.

We also adapted 3D PEG-MAL hydrogels^10^ to a 96-well plate format (Figure 1c). Briefly, fully assembled glass bottom 96-well plates were UV/ozone treated and functionalized with a solution of MPT. The MPT presents a thiol group, which reacts with the maleimide to covalently attach the 3D hydrogel to the plate. This 3D platform consisting solely of PEG and peptides is the quickest platform to fabricate: the plate functionalization, pre-polymer solution, and cell seeding can all be completed within a few hours.

### Custom-built 96-well plates with covalently bound, full-length ECM proteins and protein combinations

To create a 96-well plate system with precisely tailored integrin-binding ECM proteins, we modified commercially available glass with aminated silane (Figure 1a).^19^ We performed XPS to ensure that the silane chemistry resulted in the appropriate functional groups after each reaction step (Figure 2a). We covalently attached the cell-binding peptide RGD (GRGDSPCG) to the surface of the glass in lieu of a full-length protein due to the limitations of the XPS resolution being able to detect elements extending only a short distance from the substrate surface. Our XPS results indicated that we did not start with perfectly clean glass, demonstrated by the detection of carbon on the surface. Following APTES vapor deposition, we expected and observed an increase in carbon relative to silicon, a 3:1 ratio of carbon to nitrogen, and a 1:1 ratio of silicon to nitrogen. We observed an increase in carbon relative to silicon after the DSC reaction, with relatively unchanged ratios of oxygen and nitrogen to silicon. This indicates a substoichiometric reaction of the DSC to the surface, which could be due to imperfectly clean glass at the start, or reaction of the succinimydil carbonate with adjacent free nitrogens. Finally, with the addition of we observed an expected increase of nitrogen relative to silicon, approximately consistent with the efficacy of the DSC reaction in the third step (Figure 2b).

**Figure 2.**
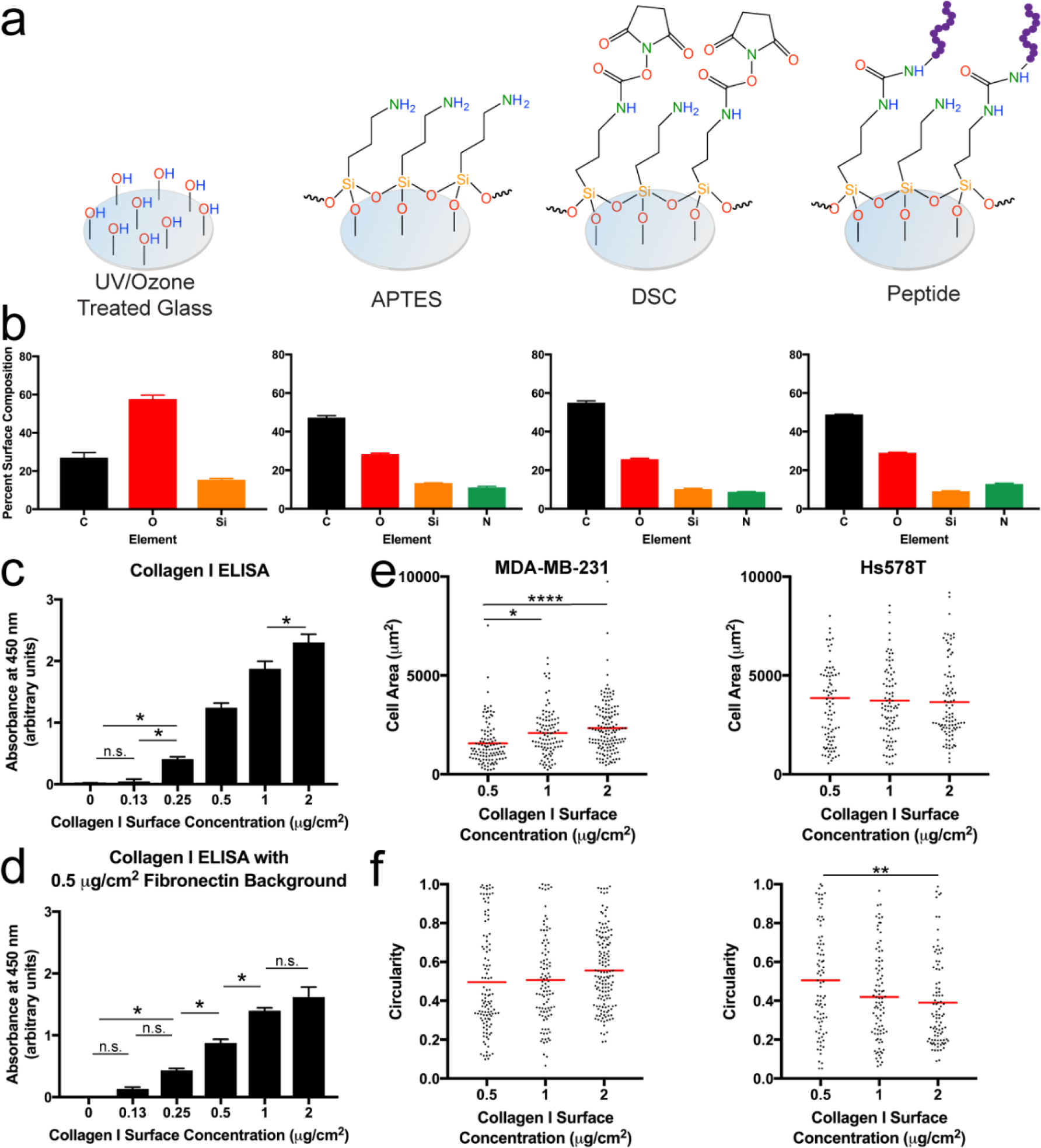
APTES-DSC chemistry allows for ECM protein coupling to glass bottom 96-well plates to control cell binding. a. Schematic of the chemical functionalization steps to create the ECM proteins attached to glass platform. b. XPS analysis of glass coverslips at each modification step. The percentage of carbon (C), oxygen (O), silicon (Si), and nitrogen (N) on the surface after each chemical step (ordered left to right). N = 2 locations per sample were used for quantification. Data are mean ± SEM. c. Modified ELISA to quantify the presence of full-length collagen I concentration on the well surfaces, N = 6 wells per condition. Data are mean ± SEM. d. Modified ELISA to quantify the presence of changing collagen I surface concentration with a constant 0.5 μg/cm^2^ fibronectin background. N = 3 wells per condition. Data are mean ± SEM. e-f. Quantification of MDA-MB-231 and Hs578T cell area (e) and cell circularity (f) as a function of collagen I surface concentrations. Individual data points are shown and the red line indicates the mean value at each condition, N ≥ 85 cells per condition. The collagen I surface concentrations on the x-axes in c-f were the expected maximum theoretical surface concentrations based on the solution concentrations that were applied to the surface. Statistical significance where p<0.05 is denoted with *, ≤0.01 with **, ≤0.001 with ***, and ≤0.0001 with ****. In Figure 2c-d statistical significance with p ≤ 0.01 are not displayed on the graph for simplicity.

To demonstrate the ability to tune ECM protein concentration and combinations, we coupled solutions containing 0-2 μg/cm^2^ of full-length collagen I to the amine-reactive glass surfaces. The surface concentrations were the expected maximum theoretical surface concentrations based on the solution concentrations. We quantified the protein coupling with a modified ELISA and observed a statistically significant increase in collagen I over the range of concentrations that we tested (Figure 2c). Real tissues are comprised of multiple proteins,^30-31^ therefore, we performed the same collagen I range quantification against a background of a constant concentration of fibronectin in the mixture. Again, we detected a statistically significant change in collagen I concentration across this range while in this ECM protein mixture (Figure 2d).

We then used this tunable glass bottom plate system to demonstrate changes in cell phenotype according to changes in collagen I surface concentrations. We quantified cell area (Figure 2e) and circularity (Figure 2f) in two breast cancer cell lines: MDA-MB-231 and Hs578T. With increasing collagen I concentration, cell area increased in the MDA-MB-231 cells, while there was no statistically significant change in Hs578T cell area (Figure 2e). MDA-MB-231 circularity did not change as a function of collagen I concentration, but there was a clear decrease in the circularity of the Hs578T cell population with increasing collagen I concentration (Figure 2f), demonstrating that this control of ECM protein has a measurable effect on typical cell behaviors.

### LAP is an efficient photoinitiator to synthesize 2D PEG-PC hydrogels in a 96-well plate format

In addition to protein composition, ^30-31^ every tissue has a specific stiffness range ^32-33^ that can be mimicked using hydrogels. We had developed a 2D PEG-PC hydrogel that is polymerized with Irgacure 2959 (Irgacure),^34^ a commonly used photoinitiator in hydrogel synthesis.^35-37^ However, this initiator could not be used in a black-walled 96-well plate due to light intensity loss. To overcome this, we used LAP,^26^ which forms free radicals more efficiently at 365 nm light than Irgacure.^26^ We made 1, 3, 6, and 10 wt% PEGDMA hydrogels with a constant 17 wt% PC. In the multi-well plates, varying the crosslinker (PEGDMA) in the hydrogels allowed for control of the effective Young’s modulus (Figure 3a). Of note, the modulus measured for hydrogels created in the 96-well plates was detectably lower than the modulus of the same condition made on glass coverslips (Figure S3a). Additionally, the bulk stiffness reported for PEG-PC hydrogels synthesized with LAP initiation was softer than what was previously reported for the hydrogels that were synthesized using Irgacure for initiation.^34^ This is likely due to weaker light transmittance in black-walled plates.^35, 37^

**Figure 3.**
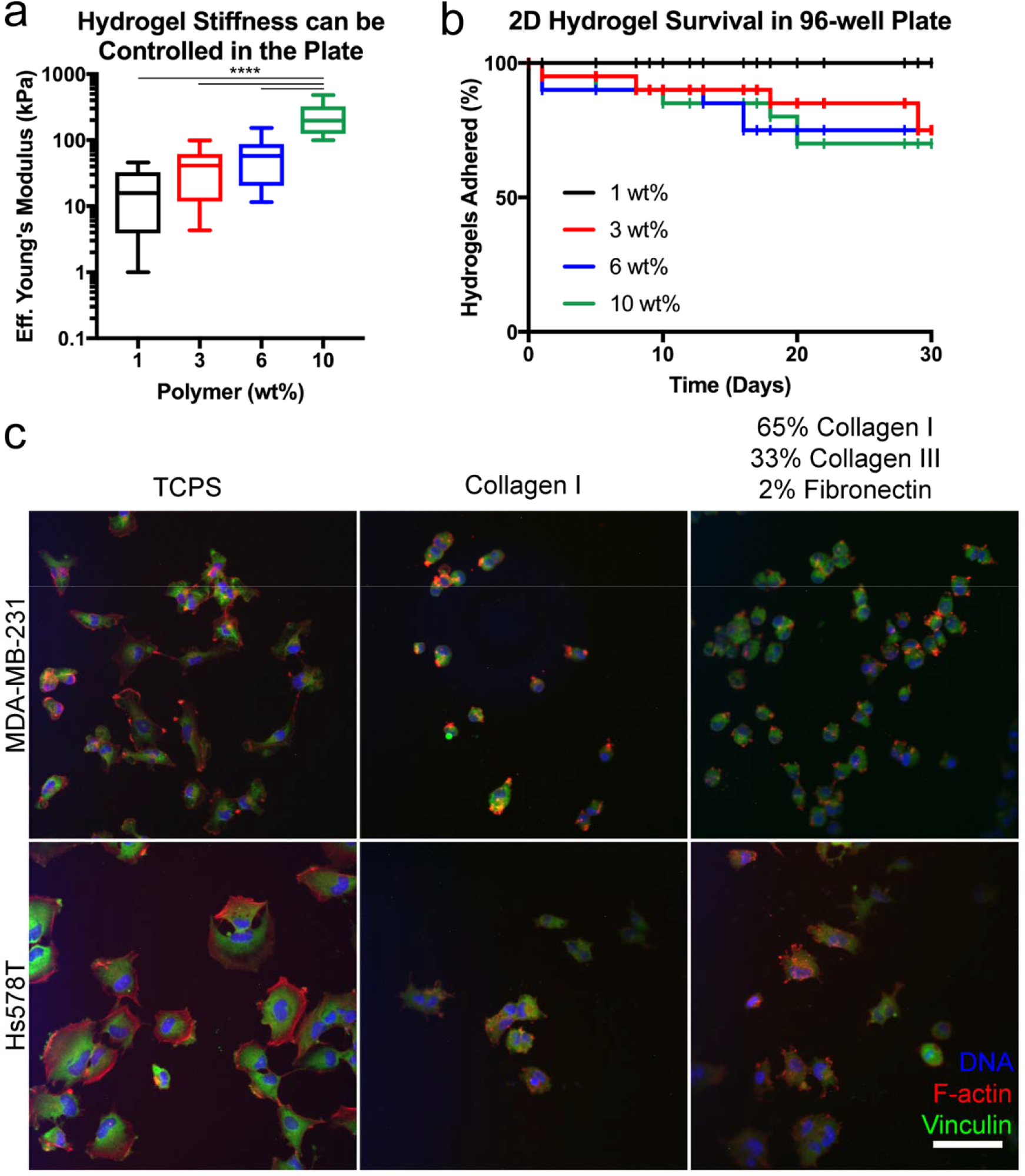
2D PEG-PC hydrogels can be synthesized in a 96-well plate with an efficient photoinitiator. a. Indentation testing of 2D PEG-PC hydrogels made at 1, 3, 6, and 10 wt% PEGDMA with a constant 17 v% solution of PC shows an increasing effective Young’s modulus with increasing PEGDMA weight percent, N = 10 hydrogels per condition. b. Percentage of the hydrogels at each condition that remain covalently attached to the MPS functionalized glass bottom 96-well plate throughout 30 days, N ≥ 6 hydrogels per condition. c. Immunofluorescence images of MDA-MB-231 and Hs578T cell lines on TCPS or PEG-PC hydrogels (10 wt% PEGDMA) coated with an expected maximum theoretical surface concentration of either 5 μg/cm^2^ collagen I or 5 μg/cm^2^ comprised of 65% collagen I, 33% collagen III, and 2% fibronectin. Cells were stained for vinculin (green), F-actin (red), and DNA in the nuclei (blue). Scale bar = 100 μm. Images are representative of N = 2 biological replicates. Statistical significance where p ≤0.0001 is denoted with ****.

We also tested the stability of the covalent attachment of these hydrogels to the glass bottom surfaces (Figure 3b). This attachment was similar comparing hydrogels attached to the glass-bottom

multi-well plates and individual coverslips (Figure S3b), suggesting that the efficacy of the silane treatment is similar between the two different glass formats.

To evaluate our ability to control protein attachment to these hydrogel surfaces, and to demonstrate a proof-of-concept study using this system to study cell behavior, MDA-MB-231 and Hs578T breast cancer cell lines were seeded on a 10 wt% hydrogel condition coupled with an expected maximum theoretical surface concentration of 5 μg/cm collagen I alone or a “collagen-rich” mixture (65% collagen I, 33% collagen III, & 2% fibronectin).^6^ After 24 hours, we fixed and stained the cells (Figure 3c). Both cell lines spread out less on the soft hydrogels compared to TCPS, confirming the results of many other groups.^6, 34, 38–41^ Cells were unable to spread out on 2D hydrogels that were not functionalized with protein (Figure S3c-d). Comparing the 2D hydrogels by qualitative visual inspection, we observed that more cells could adhere to the “collagen-rich” substrate over the collagen I only substrate for both MDA-MB-231 and Hs578T cell lines. This result is likely due to the variety of integrin-binding sites provided by the protein mixture. Largely in agreement with our previous work on PEG-PC gels, we saw very little evidence of punctate focal adhesions in any of the conditions tested, which appears to be a general feature of these breast cancer cell lines.^34^

### PEG-Maleimide hydrogels with encapsulated cells in 3D were created with liquid handling robotics

Cells experience a 3D microenvironment *in vivo* and we adapted our 3D PEG-MAL hydrogel^10^ to 96-well plates via liquid handling robotics to increase the throughput of this platform for studying cells in 3D. As in our PEG-PC system, we could control the effective Young’s modulus of the hydrogels within the well plates by tuning the polymer weight percent (Figure 4a). Also, there was no significant difference between the stiffnesses measured for hydrogels synthesized on coverslips or in multi-well plates (Figure S4a), and the values reported for these hydrogels is consistent with previous reports.^27^ This result was expected since this reaction does not require UV light irraditation.^27^

**Figure 4.**
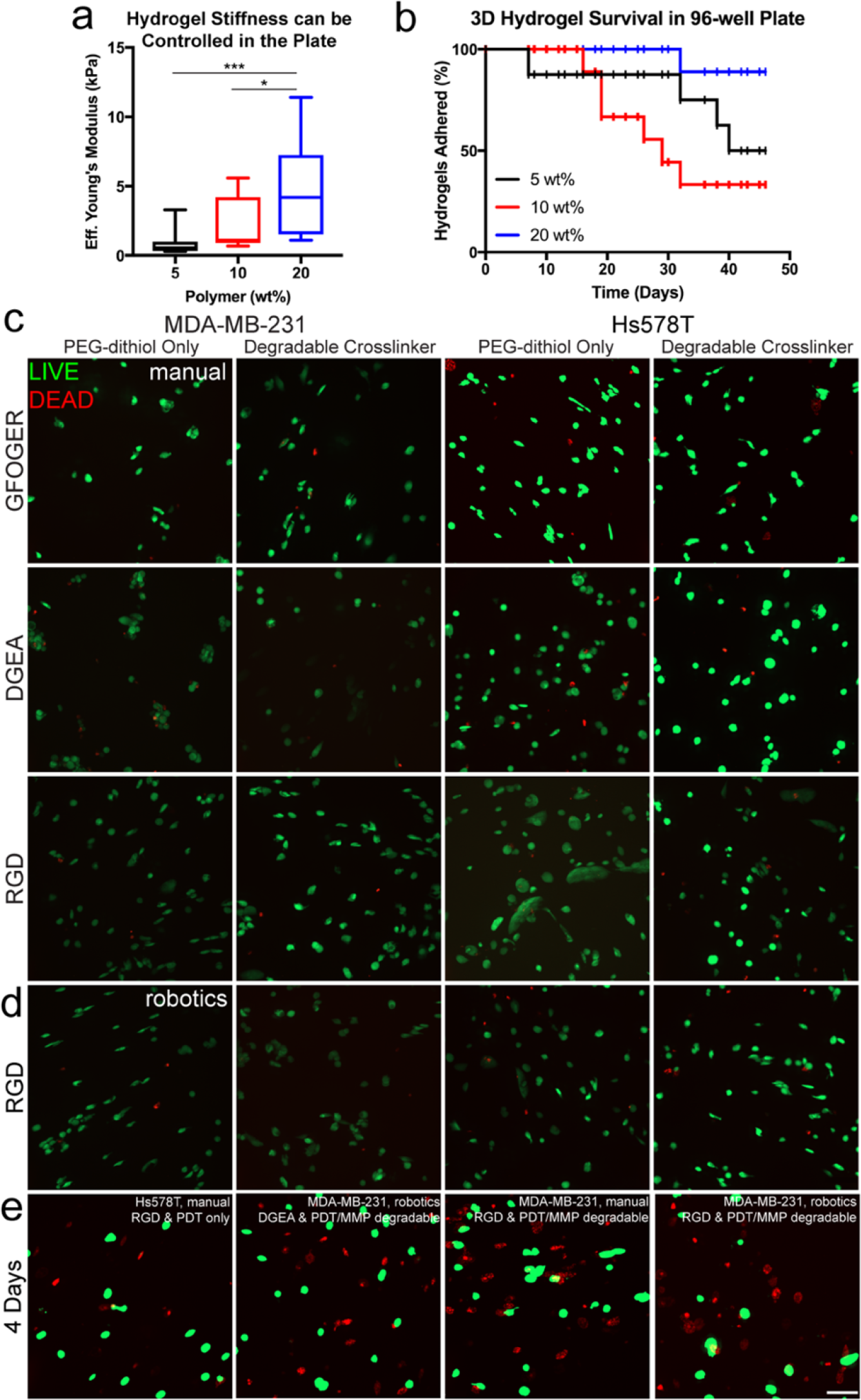
Silane functionalization allows for the synthesis of 3D PEG-MAL hydrogels in a 96-well plate format. a. Indentation testing of 5, 10, and 20 wt% 3D 4-arm PEG-MAL hydrogels crosslinked with linear PEG-dithiol measured the effective Young’s modulus of hydrogels in a 96-well plate, N ≥ 11 hydrogels per condition. b. Long-term 3D PEG-MAL hydrogel covalent attachment to the glass bottom 96-well plates, which were treated with MPT, N ≥ 6 hydrogels per condition. c. LIVE (green)/DEAD (red) imaging of MDA-MB-231 and Hs578T cells one day post-encapsulation with manual pipetting in the 3D hydrogel with 2 mM RGD, DGEA, or GFOGER celladhesion peptides and crosslinked with PEG-dithiol or PEG-dithiol plus a Pan-MMP degradable sequence (indicated as degradable crosslinker). d. LIVE (green)/DEAD (red) imaging of MDA-MB-231 and Hs578T cells one day after encapsulation with liquid handling robotics in the 3D hydrogel with 2 mM RGD cell-adhesion peptide and crosslinked with PEG-dithiol or PEG-dithiol plus a Pan-MMP degradable sequence. Images are representative from N ≥ 3 biological replicates. e. LIVE (green)/DEAD (red) imaging of MDA-MB-231 and Hs578T cells four days post-encapsulation at selected conditions. Scale bar = 100 μm. Statistical significance where p<0.05 is denoted with * and ≤0.001 with ***.

During gelation, the 3D PEG-MAL hydrogels covalently attached to the glass bottom plates by reacting to the thiol group we silane-functionalized to the surface. Most of the 3D PEG-MAL hydrogels remained adhered to the well surface for more than 30 days (Figure 4b). This was also observed for the 2D PEG-PC hydrogels (Figure 3b), and this stability was dependent on the polymer weight percent, likely due to the number of available maleimide groups to react with the silane-functionalized surface and how much the different Hydrogels swell with the well (Figure 4b). it was not possible io synmesize inese 3D PEG-MAL hydrogels in plates that were not silane treated, because the hydrogels wicked to the sides of the wells. When we synthesized the hydrogels on silane treated coverslips, they adhered for more than 100 days (Figure S4b-c).

As a proof-of-concept study, we encapsulated MDA-MB-231 or Hs578T cells in 3D PEG-MAL hydrogels with both manual pipetting (Figure 4c,e) and liquid handling robotics (Figure 4d-e). We chose three peptide sequences, RGD, GFOGER, and DGEA, that represent cell-adhesion sites in collagen I (GFOGER & DGEA)^42-44^ and fibronectin/vitronectin/osteopontin (RGD).^43,45^ Additionally, an enzyme-degradable crosslinker (Pan-MMP, GPQG↓IWGQ) was incorporated into some of the hydrogels to allow the cells to degrade and move through the matrix. Cell viability was measured at one and four days post-encapsulation via LIVE/DEAD staining (Figure 4c-e, S5). The cells encapsulated in hydrogels without cell-adhesion peptides were still mostly viable one day post-encapsulation, but at four days post-encapsulation the cell viability was clearly much lower as expected since these are adherent cell lines (Figure S5a). Cells were over 80% viable within the hydrogels one day post-encapsulation regardless of whether the hydrogels were made manually or via liquid handling robotics (Figure 4c-d). The cells encapsulated in the Pan-MMP degradable hydrogel had higher viability (>60%) at day four than the PDT only crosslinking condition (<60%, Figure S5b). We speculate that this increase in cell viability is due to the degradable sequences provided, which facilitated higher cell spreading, a possible explanation in the difference in viability. Overall, these results indicate that the PEG-MAL hydrogel system is adaptable to automated liquid handling robotics, and the incorporation of degradable sequences in the matrix improves cell viability.

## DISCUSSION

Many groups have developed biomaterial cell culture platforms and suggested that future applications could include drug screening.^4-5,46-47^ These materials could have significant impact on improving the data obtained by *in vitro* cell studies, if adapted to current high-throughput screening technologies. Towards this goal, we adapted our lab’s existing biomaterial technologies to 96-well plates and semi-automated liquid handling robotics. Our ability to adapt these materials from fully manual fabrication methods to semi-automated fabrication in 96-welll plates demonstrated that these materials are adaptable to liquid handling robotics. This is an important step for eventual implementation in an industrial setting, or even lab-scale screening studies. In our case, the liquid handling robotics is shared equipment, specifically configured for 250 μL syringe pumps for pipetting. Although not possible for us to do in this study, modifying the syringe pumps can increase the pipetting capacity of the system. For this reason, we limited the use of the robotics for pipetting repeated volumes greater than 100 μL. This fact meant that in our study, the semi-automated liquid handling robotics saved only a moderate amount of time relative to manual pipetting when making a few plates at a time. However, if more than 10 plates are needed for a study, the robotic workstation becomes a time-saver, and this savings scales upward with the number of plates. Protocols to use liquid handling robotics have been established for TCPS plates, but the incorporation of a variety of different types of biomaterials is a contribution we make here as a mechanism to enable higher-throughput screening with biomaterials *in vitro*.

Since the importance of cell culture dimensionality has been increasingly evident,^48^ a tunable, reproducible, and easily fabricated high-throughput 3D *in vitro* cell culture platform is a valuable tool for drug screening.^48-49^ The Lutolf group has demonstrated the use of 3D hydrogels in a microarray made using automated liquid handling robotics to test many microenvironmental conditions on the regulation of mouse embryonic stem cell self-renewal.^50^ This study was a valuable demonstration of the utility of biomaterials in a high-throughput format and it was an important contribution towards using synthetic materials for large-scale *in vitro* screens. In our previous 3D PEG-MAL hydrogels drug screening study,^10^ we used 10 μL hydrogel volumes to avoid diffusion limitations, and in 96-well plates this volume wicked to the edge of the wells, which limited our study 48-well plates. Here, we overcame this challenge by functionalizing glass well surfaces with thiol-terminated silane to covalently attach the 3D hydrogels to the surface. The stiffness of the hydrogels in the plate was tunable (Figure 4a), and these hydrogels also stuck reliably to the surface of the plate for over 30 days (Figure 4b), which is critical for long-term cell culture studies. These PEG-maleimide-based hydrogels are very compliant and are appropriate for mimicking soft tissues. The silane chemistry could be used to attach hydrogels up to 100 days (Figure S4c), indicating that this surface preparation can be done well in advance of hydrogel synthesis.

The chemistry of PEG-MAL hydrogels allows for gelation at neutral pH,^27^ and the encapsulated cells were not exposed to UV light, as is common in other systems.^9,11^ The PEG-based hydrogel system allows the specific incorporation of cell-binding and cell-degradable functional groups, which is not possible with protein-based gels such as collagen^51^ or Matrigel.^52^ Moreover, these protein based gels have high amounts of batch-to-batch variability,^53^ which makes the reproducibility of results difficult in large scale drug screening assays. Since our synthetic 3D model only requires two components: 1) PEG, which allows for tuning the hydrogel stiffness, and 2) peptides, which allow for cell attachment to the matrix and hydrogel degradation, it is more reproducible than protein-based hydrogels where there is known lot-to-lot variability. To increase the throughput of this platform, we synthesized the hydrogels with semi-automated liquid handling robotics. After isolating the appropriate cells and preparing the hydrogel precursor solutions, one plate of these hydrogels can be synthesized in less than five minutes with liquid handling robotics, where manual pipetting takes about 20 minutes. Further, as an extension of the study of ECM proteins attached to glass, we incorporated peptides representative of collagen I (GFOGER & DGEA) or fibronectin/vitronectin/osteopontin (RGD) binding sequences and a Pan-MMP degradable sequence. The MDA-MB-231 and Hs578T cells maintained over 80% viability for all conditions one day post-seeding (Figure 4c-d, S5b). Cell viability was also maintained with the use of liquid handling robotics (Figure 4d-e). Looking specifically at Hs578T cells encapsulated with RGD, there was lower survival on day four than on day one. This may be due to a need for additional integrin-binding sites other than RGD or an increase in total binding sites for long-term survival in the hydrogel. This result was important for scaling up our 3D PEG-MAL hydrogel system for high-throughput applications.

For situations where full-length proteins are essential, we also demonstrated a silane-based approach to create multi-well plate 2D hydrogels. This also allows users to capture stiffness when 3D geometry is unnecessary to answer the biological question at hand. Others have developed 2D hydrogel platforms,^7, 20^ which produce perfectly flat hydrogels, but they are complicated to fabricate^7^ or require a plate insert,^20^ limiting the number of plates that can be made at once. Microarrays with multiple 2D hydrogel stiffnesses and protein binding surfaces have been developed to demonstrate that both factors impact mesenchymal stem cell behavior.^3^ Hydrogels with incorporated nanofibers to control the spatial arrangement of biochemical signals have also been developed,^54^ but this type of system is complicated to fabricate. Our original 96-well plate 2D PEG-PC hydrogel system design^6^ was hindered for scale up because gelation required an oxygen-free environment. Irgacure is a photoinitiator that is commonly used for initiating hydrogel polymerization, ^35-37^ and it was used for 2D PEG-PC hydrogels when they were first developed for coverslips.^34^ Irgacure is not a very efficient photoinitiator^26^ because it does not absorb light readily at 365 nm,^35^ but this wavelength is used in other platforms that include cell encapsulation to limit damage to the cells. ^35,55-56^ Since the pre-polymer solution resides deep within the wells of black-walled plates, the efficacy of Irgacure was particularly problematic, and therefore it was not possible to form PEG-PC hydrogels in 96-well plates. Since LAP is more efficient than Irgacure, it was possible to make the PEG-PC hydrogels in 96-well plates using this photoinitiator. This allows the use of liquid handling robotics for pipetting the pre-polymer solution without initiating polymerization of hydrogels in the pipette tips.

Using LAP, we could control hydrogel stiffness within the plate across three orders of magnitude, enabling this material to mimic most soft and elastic tissues (Figure 3a).^33-34^ Also, the silane treatment allowed for the long-term adherence of the hydrogels to the well surface (Figure 3b). Both MDA-MB-231 and Hs578T breast cancer cell spreading was mechanosensitive, in agreement with observations by us and many other groups.^6,34,38-41^ Specific to our study, both cell lines adhered and spread more on the “collagen-rich” mixture than with collagen I alone (Figure 3c), perhaps due to the additional integrin binding sites provided. Both MDA-MB-231 and Hs578T breast cancer cell lines express high levels of β_1_ integrin,^57^ which pairs with multiple integrin α subunits to bind to collagen I and fibronectin, possibly explaining the result we observed.

To isolate cell-ECM binding, we developed a third form of silane-treated glass-bottom 96-well plate. Previous work with this platform had been limited to a 24-well plate,^19^ where treated glass coverslips were prepared and inserted by hand. Unlike the 2D and 3D hydrogel platforms, the chemical functionalization to the glass cannot be performed in a fully assembled glass-bottom well plate due to the incompatibility of polystyrene and some of the solvents. Therefore, the custom 96-well plate format was created by first modifying one well plate-sized piece of glass and attaching it to a commercially available bottomless 96-well plate with a medical-grade adhesive. The benefits of using of a 96-well plate for covalently attaching ECM proteins to glass include both a reduction in costly materials, such as full-length proteins, and a savings in time to prepare the platform, particularly since we chemically modified and attached a single piece of glass instead of individual glass coverslips. This approach presents a feasible method for scaling up to 384 or 1,536 well plates, which are also commonly used in industrial applications. If this approach is adopted by industry, the reproducibility of the chemistry is essential for quality control. We noted from XPS that the reaction stoichiometry was not 100% efficient (Figure 2a-b). This could be due to the starting glass not being completely pure/clean or some amount of APTES layer peeling typical of multilayer silane formation ^25^ In an attempt to maximize silane monolayers *vs* multilayers, our silane deposition was done in the vapor phase, but multilayer silanes are still possible. Regardless, we showed that the surface concentration of protein in each well can be carefully controlled (Figure 2c-d), and these 2D biomaterial platforms could be used as a straightforward way to study tumor progression where collagen density increases and ECM composition changes.^58^ We observed an overall trend of increasing cell area in MDA-MB-231 cells with increasing collagen I surface concentration (Figure 2e), alongside a decrease in the circularity of the Hs578T cells (Figure 2f). These types of studies may help identify a certain threshold of ligand density where efficacy of a compound is greatly enhanced or reduced.

The use of synthetic biomaterials allows for the capture of both biochemical and mechanical cues experienced by cells *in vivo*. These rationally designed *in vitro* models are beneficial because components of the microenvironment can be controlled individually, which is an impossible task *in vivo.* The likely application of interest for each platform will be dependent on the user’s end goal. For example, the ECM proteins attached to glass and 2D PEG-PC hydrogels incorporate full-length proteins, whereas the 3D PEG-MAL hydrogels only include peptides representative of integrin-binding sequences to specific proteins. Since the structure of the protein can impact integrin-binding,^59-60^ it is an important consideration in biomaterial choice. Our goal was to present mechanisms to use each of these platforms in 96-well plates with semi-automated liquid handling robotics to enable high-throughput applications in the future. Specific features included in each, such as protein versus peptide, stiffness, and geometry will guide their exact applications in individual labs or companies.

## CONCLUSIONS

We have adapted three unique, but complementary biomaterial platforms to 96-well plates and used semi-automated liquid handling robotics to enable high-throughput screening applications. Each platform was modified from existing technology to reduce material requirements and fabrication time per condition. Specifically, our 2D hydrogels can be synthesized with a photoinitiator in a 96-well plate in the presence of oxygen, our 3D hydrogels synthesis can be semi-automated to reduce the time invested per plate by four-fold, and our custom 96-well plate design enables screening on controlled glass substrate compositions. While more complex and expensive than traditional TCPS, these systems include specific microenvironmental features that could be key to better predict *in vivo* cell behavior, either for applications in preclinical drug screening or as better tools for hypothesis test-beds.

## AUTHOR INFORMATION

### Author Contributions

EAB and SRP designed the experiments. AMY performed the saline stability experiment and EAB, LEJ, and MFG performed and analyzed all of the other experiments. SRP supervised the study. EAB and SRP wrote the manuscript and MFG and LEJ contributed to editing.

## Notes

The authors declare no competing financial interests.

## ACKNOWLEDGEMENTS

The authors are grateful to Dr. Alfred Crosby and Dr. Jungwoo Lee for use of equipment and Jun-Goo Kwak for technical assistance. Dr. Thomas McCarthy allowed for the use of XPS equipment and Jack Hirsch performed the analysis. ARcare 90106 was donated by Adhesives Research. This work was funded by a grant from the NIH (1DP2CA186573-01) and a CAREER grant from the NSF (1454806) to SRP. SRP is a Pew Biomedical Scholar supported by the Pew Charitable Trusts. SRP was supported by a Barry and Afsaneh Siadat Career Development Award. EAB was partially supported by National Research Service Award T32 GM008515 from the National Institutes of Health. MFG was partially supported by the Barry and Afsaneh Siadat Career Development Award.

## SUPPORTING INFORMATION

### Supplemental Materials and Methods

DGEA synthesis

**Figure S1:** Custom glass bottom 96-well plates can be made with commercially available materials.

**Figure S2:** XPS survey traces for the chemical modification steps to prepare ECM proteins attached to glass.

**Figure S3:** 2D PEG-PC hydrogels can also be made on coverslips and cells do not attach to hydrogels without protein functionalization.

**Figure S4:** 3D PEG-MAL hydrogels can also be made on coverslips.

**Figure S5:** Cell viability in 3D PEG-MAL hydrogels.

## REFERENCES

1. Gjorevski, N.; Sachs, N.; Manfrin, A.; Giger, S.; Bragina, M. E.; Ordóñez-Morán, P.; Clevers, H.; Lutolf, M. P., Designer matrices for intestinal stem cell and organoid culture. Nature 2016, 539 (7630), 560-564. DOI: 10.1038/nature20168.

2. Banks, J. M.; Harley, B. A. C.; Bailey, R. C., Tunable, Photoreactive Hydrogel System To Probe Synergies between Mechanical and Biomolecular Cues on Adipose-Derived Mesenchymal Stem Cell Differentiation. ACS Biomater. Sci. Eng. 2015, 1 (8), 718-725. DOI: 10.1021/acsbiomaterials.5b00196.

3. Gobaa, S.; Hoehnel, S.; Lutolf, M. P., Substrate elasticity modulates the responsiveness of mesenchymal stem cells to commitment cues. Integr. Biol. 2015, 7 (10), 1135-1142. DOI: 10.1039/C4IB00176A.

4. Casey, J.; Yue, X.; Nguyen, T. D.; Acun, A.; Zellmer, V. R.; Zhang, S.; Zorlutuna, P., 3D hydrogel-based microwell arrays as a tumor microenvironment model to study breast cancer growth. Biomed. Mater. 2017, 12 (2), 1-12. DOI: 10.1088/1748-605X/aa5d5c.

5. Bray, L. J.; Binner, M.; Holzheu, A.; Friedrichs, J.; Freudenberg, U.; Hutmacher, D. W.; Werner, C., Biomaterials Multi-parametric hydrogels support 3D in vitro bioengineered microenvironment models of tumour angiogenesis. Biomaterials 2015, 53, 609-620. DOI: 10.1016/j.biomaterials.2015.02.124.

6. Nguyen, T. V.; Sleiman, M.; Moriarty, T.; Herrick, W. G.; Peyton, S. R., Sorafenib resistance and JNK signaling in carcinoma during extracellular matrix stiffening. Biomaterials 2014, 35 (22), 5749-5759. DOI: 10.1016/j.biomaterials.2014.03.058.

7. Zustiak, S.; Nossal, R.; Sackett, D. L., Multiwell stiffness assay for the study of cell responsiveness to cytotoxic drugs. Biotechnol. Bioeng. 2014, 111 (2), 396-403. DOI: 10.1002/bit.25097.

8. Barney, L. E.; Jansen, L. E.; Polio, S. R.; Galarza, S.; Lynch, M. E.; Peyton, S. R., The predictive link between matrix and metastasis. Curr. Opin. Chem. Eng. 2016, 11, 85-93. DOI: 10.1016/j.coche.2016.01.001.

9. Chen, J.-W. E.; Pedron, S.; Harley, B. A. C., The Combined Influence of Hydrogel Stiffness and Matrix-Bound Hyaluronic Acid Content on Glioblastoma Invasion. Macromol. Biosci. 2017, 17 (8), 718-725. DOI: 10.1002/mabi.201700018.

10. Gencoglu, M. F.; Barney, L. E.; Hall, C. L.; Brooks, E. A.; Schwartz, A. D.; Corbett, D. C.; Stevens, K.; Peyton, S. R., Comparative study of multicellular tumor spheroid formation methods and implications for drug screening. ACS Biomater. Sci. Eng. 2017. DOI: 10.1021/acsbiomaterials.7b00069.

11. Mabry, K. M.; Schroeder, M. E.; Payne, S. Z.; Anseth, K. S., Three-Dimensional High-Throughput Cell Encapsulation Platform to Study Changes in Cell-Matrix Interactions. ACS Appl. Mater. Interfaces 2016, 8 (34), 21914-21922. DOI: 10.1021/acsami.5b11359.

12. Sawicki, L. A.; Kloxin, A. M., Light-mediated Formation and Patterning of Hydrogels for Cell Culture Applications. J. Visualized Exp. 2016, (115), 1-10. DOI: 10.3791/54462.

13. Torisawa, Y.-s.; Spina, C. S.; Mammoto, T.; Mammoto, A.; Weaver, J. C.; Tat, T.; Collins, J. J.; Ingber, D. E., Bone marrow–on–a–chip replicates hematopoietic niche physiology in vitro. Nat. Methods 2014, 11 (6), 663-669. DOI: 10.1038/nmeth.2938.

14. Li, C. Y.; Stevens, K. R.; Schwartz, R. E.; Alejandro, B. S.; Huang, J. H.; Bhatia, S. N., Micropatterned cell–cell interactions enable functional encapsulation of primary hepatocytes in hydrogel microtissues. Tissue Eng., Part A 2014, 20 (15 and 16), 2200-2212. DOI: 10.1089/ten.tea.2013.0667.

15. Montanez-Saur, S. I.; Sung, K. E.; Berthier, E.; Beebe, D. J., Enabling screening in 3D microenvironments: probing matrix and stromal effects on the morphology and proliferation of T47D breast carcinoma cells. Integr. Biol. 2013, 5 (3), 631-640. DOI: 10.1039/c3ib20225a.

16. Jeon, J. S.; Bersini, S.; Gilardi, M.; Dubini, G.; Charest, J. L.; Moretti, M.; Kamm, R. D., Human 3D vascularized organotypic microfluidic assays to study breast cancer cell extravasation. Proc. Natl. Acad. Sci. U. S. A. 2015, 112 (1), 214-219. DOI: 10.1073/pnas.1501426112.

17. Benam, K. H.; Villenave, R.; Lucchesi, C.; Varone, A.; Hubeau, C.; Lee, H.-H.; Alves, S. E.; Salmon, M.; Ferrante, T. C.; Weaver, J. C.; Bahinski, A.; Hamilton, G. A.; Ingber, D. E., Small airway-on-a-chip enables analysis of human lung inflammation and drug responses in vitro. Nat. Methods 2016, 13 (2), 151-157. DOI: 10.1038/nmeth.3697.

18. Fan, Y.; Nguyen, D. T.; Akay, Y.; Xu, F.; Akay, M., Engineering a Brain Cancer Chip for High-throughput Drug Screening. Sci. Rep. 2016, 6, 25062. DOI: 10.1038/srep25062.

19. Barney, L. E.; Dandley, E. C.; Jansen, L. E.; Reich, N. G.; Mercurio, A. M.; Peyton, S. R., A cell-ECM screening method to predict breast cancer metastasis. Integr. Biol. 2015, 7, 198-212. DOI: 10.1039/C4IB00218K.

20. Mih, J. D.; Sharif, A. S.; Liu, F.; Marinkovic, A.; Symer, M. M.; Tschumperlin, D. J., A Multiwell Platform for Studying Stiffness-Dependent Cell Biology. PLoS One 2011, 6 (5), e19929. DOI: 10.1371/journal.pone.0019929.

21. Tokuda, E. Y.; Jones, C. E.; Anseth, K. S., PEG-peptide hydrogels reveal differential effects of matrix microenvironmental cues on melanoma drug sensitivity. Integr. Biol. 2017, 9 (1), 76-87. DOI: 10.1039/C6IB00229C.

22. Hay, M.; Thomas, D. W.; Craighead, J. L.; Economides, C.; Rosenthal, J., Clinical development success rates for investigational drugs. Nat. Biotechnol. 2014, 32 (1), 40-51. DOI: 10.1038/nbt.2786.

23. Hertzberg, R. P.; Pope, A. J., High-throughput screening: new technology for the 21st century. Curr. Opin. Chem. Biol. 2000, 4 (4), 445-451. DOI: 10.1016/S1367-5931(00)00110-1.

24. Harkness, T.; McNulty, J. D.; Prestil, R.; Seymour, S. K.; Klann, T.; Murrell, M.; Ashton, R. S.; Saha, K., High-content imaging with micropatterned multiwell plates reveals influence of cell geometry and cytoskeleton on chromatin dynamics. Biotechnol. J. 2015, 10 (10), 1555-1567. DOI: 10.1002/biot.201400756.

25. Zhu, M.; Lerum, M. Z.; Chen, W., How To Prepare Reproducible, Homogeneous, and Hydrolytically Stable Aminosilane-Derived Layers on Silica. Langmuir 2012, 28, 416-423. DOI: 10.1021/la203638g.

26. Fairbanks, B. D.; Schwartz, M. P.; Bowman, C. N.; Anseth, K. S., Biomaterials Photoinitiated polymerization of PEG-diacrylate with lithium phenyl-2,4,6-trimethylbenzoylphosphinate: polymerization rate and cytocompatibility. Biomaterials 2009, 30 (35), 6702-6707. DOI: 10.1016/j.biomaterials.2009.08.055.

27. Phelps, E. A.; Enemchukwu, N. O.; Fiore, V. F.; Sy, J. C.; Murthy, N.; Sulchek, T. A.; Barker, T. H.; García, A. J., Maleimide cross-linked bioactive PEG hydrogel exhibits improved reaction kinetics and cross-linking for cell encapsulation and in situ delivery. Adv. Mater. 2012, 24 (1), 64-70. DOI: 10.1002/adma.201103574.

28. Chan, E. P.; Smith, E. J.; Hayward, R. C.; Crosby, A. J., Surface Wrinkles for Smart Adhesion. Adv. Mater. 2008, 20 (4), 711-716. DOI: 10.1002/adma.200701530.

29. Hutchens, S. B.; Crosby, A. J., Soft-solid deformation mechanics at the tip of an embedded needle. Soft Matter 2014, 10, 3679-3684. DOI: 10.1039/C3SM52689E.

30. Naba, A.; Clauser, K. R.; Hoersch, S.; Liu, H.; Carr, S. A.; Hynes, R. O., The Matrisome: In Silico Definition and In Vivo Characterization by Proteomics of Normal and Tumor Extracellular Matrices. Mol. Cell. Proteomics 2012, 11 (4). DOI: 10.1074/mcp.M111.014647.

31. Uhlén, M.; Fagerberg, L.; Hallström, B. M.; Lindskog, C.; Oksvold, P.; Mardinoglu, A.; Sivertsson, Å.; Kampf, C.; Sjöstedt, E.; Asplund, A.; Olsson, I.; Edlund, K.; Lundberg, E.; Navani, S.; Szigyarto, C. A.-k.; Odeberg, J.; Djureinovic, D.; Takanen, J. O.; Hober, S.; Alm, T.; Edqvist, P.-h.; Berling, H.; Tegel, H.; Mulder, J.; Rockberg, J.; Nilsson, P.; Schwenk, J. M.; Hamsten, M.; Feilitzen, K. V.; Forsberg, M.; Persson, L.; Johansson, F.; Zwahlen, M.; Heijne, G. V.; Nielsen, J.; Pontén, F., Tissue-based map of the human proteome. Science 2015, 347 (6220). DOI: 10.1126/science.1260419.

32. Jansen, L. E.; Birch, N. P.; Schiffman, J. D.; Crosby, A. J.; Peyton, S. R., Mechanics of Intact Bone Marrow. J. Mech. Behav. Biomed. Mater. 2015, 50, 299-307. DOI: 10.1016/j.jmbbm.2015.06.023.

33. Handorf, A. M.; Zhou, Y.; Halanski, M. A.; Li, W.-J., Tissue stiffness dictates development, homeostasis, and disease progression. Organogenesis 2015, 11 (1), 1-15. DOI: 10.1080/15476278.2015.1019687.

34. Herrick, W. G.; Nguyen, T. V.; Sleiman, M.; McRae, S.; Emrick, T. S.; Peyton, S. R., PEG-Phosphorylcholine Hydrogels As Tunable and Versatile Platforms for Mechanobiology. Biomacromolecules 2013, 14 (7), 2294-2304. DOI: 10.1021/bm400418g.

35. Bryant, S. J.; Nuttelman, C. R.; Anseth, K. S., Cytocompatibility of UV and visible light photoinitiating systems on cultured NIH/3T3 fibroblasts in vitro. J. Biomater. Sci., Polym. Ed. 2000, 11 (5), 439-457. DOI: 10.1163/156856200743805.

36. Fedorovich, N. E.; Oudshoorn, M. H.; Geemen, D. V.; Hennink, W. E.; Alblas, J.; Dhert, W. J. A., The effect of photopolymerization on stem cells embedded in hydrogels. Biomaterials 2009, 30 (3), 344-353. DOI: 10.1016/j.biomaterials.2008.09.037.

37. Mironi-Harpaz, I.; Yingquan, D.; Venkatraman, S.; Seliktar, D., Photopolymerization of cell-encapsulating hydrogels: crosslinking efficiency versus cytotoxicity. Acta Biomater. 2012, 8 (5), 1838-1848. DOI: 10.1016/j.actbio.2011.12.034.

38. Peyton, S. R.; Raub, C. B.; Keschrumrus, V. P.; Putnam, A. J., The use of poly (ethylene glycol) hydrogels to investigate the impact of ECM chemistry and mechanics on smooth muscle cells. Biomaterials 2006, 27 (28), 4881-4893. DOI: 10.1016/j.biomaterials.2006.05.012.

39. Peyton, S. R.; Kim, P. D.; Ghajar, C. M.; Seliktar, D.; Putnam, A. J., The effects of matrix stiffness and RhoA on the phenotypic plasticity of smooth muscle cells in a 3-D biosynthetic hydrogel system. Biomaterials 2008, 29 (17), 2597-2607. DOI: 10.1016/j.biomaterials.2008.02.005.

40. Marklein, R. A.; Burdick, J. A., Spatially controlled hydrogel mechanics to modulate stem cell interactions. Soft Matter 2010, 6 (1), 136-143. DOI: 10.1039/b916933d.

41. Weng, S.; Fu, J., Synergistic regulation of cell function by matrix rigidity and adhesive pattern. Biomaterials 2011, 32 (36), 9584-9593. DOI: 10.1016/j.biomaterials.2011.09.006.

42. Zhang, W.-M.; Käpylä, J.; Puranen, J. S.; Knight, C. G.; Tiger, C.-F.; Pentikäinen, O. T.; Johnson, M. S.; Farndale, R. W.; Heino, J.; Gullberg, D., α11β1 Integrin Recognizes the GFOGER Sequence in Interstitial Collagens. J. Biol. Chem. 2003, 278 (9), 7270-7277. DOI: 10.1074/jbc.M210313200.

43. Yamada, K. M., Adhesive Recognition Sequences. J. Biol. Chem. 1991, 266 (20), 12809-12812.

44. Staatz, W. D.; Fok, K. F.; Zutter, M. M.; Adams, S. P.; Rodriguez, B. A.; Santoro, S. A., Identification of a Tetrapeptide Recognition Sequence for the alpha2beta1 Integrin in Collagen. J. Biol. Chem. 1991, 266 (12), 7363-7367.

45. Ruoslahti, E., RGD and other recognition sequences for integrins. Annu. Rev. Cell Dev. Biol. 1996, 12, 697-715. DOI: 10.1146/annurev.cellbio.12.1.697.

46. Pradhan, S.; Hassani, I.; Seeto, W. J.; Lipke, E. A., PEG-fibrinogen hydrogels for threedimensional breast cancer cell culture. J. Biomed. Mater. Res., Part A 2016, 105 (1), 236-252. DOI: 10.1002/jbm.a.35899.

47. Seo, J.; Shin, J.-y.; Leijten, J.; Jeon, O., Biomaterials High-throughput approaches for screening and analysis of cell behaviors. Biomaterials 2018, 153, 85-101. DOI: 10.1016/j.biomaterials.2017.06.022.

48. Baker, B. M.; Chen, C. S., Deconstructing the third dimension: how 3D culture microenvironments alter cellular cues. J. Cell Sci. 2012, 125 (Pt13), 3015-3024. DOI: 10.1242/jcs.079509.

49. Edmondson, R.; Broglie, J. J.; Adcock, A. F.; Yang, L., Three-Dimensional Cell Culture Systems and Their Applications in Drug Discovery and Cell-Based Biosensors. Assay Drug Dev. Technol. 2014, 12 (4), 207-218. DOI: 10.1089/adt.2014.573.

50. Ranga, A.; Gobaa, S.; Okawa, Y.; Mosiewicz, K.; Negro, A.; Lutolf, M. P., 3D niche microarrays for systems-level analyses of cell fate. Nature Communications 2014, 5, 1-10. DOI: 10.1038/ncomms5324.

51. Szot, C. S.; Buchanan, C. F.; Freeman, J. W.; Rylander, M. N., 3D in vitro bioengineered tumors based on collagen I hydrogels. Biomaterials 2011, 32 (31), 7905-7912. DOI: 10.1016/j.biomaterials.2011.07.001.

52. Kleinman, H. K.; Martin, G. R., Matrigel: Basement membrane matrix with biological activity. Semin. Cancer Biol. 2005, 15 (5), 378-386. DOI: 10.1016/j.semcancer.2005.05.004.

53. Hughes, C. S.; Postovit, L. M.; Lajoie, G. A., Matrigel: a complex protein mixture required for optimal growth of cell culture. Proteomics 2010, 10 (9), 1886-1890. DOI: 10.1002/pmic.200900758.

54. Wade, R. J.; Bassin, E. J.; Gramlich, W. M.; Burdick, J. A., Nanofibrous Hydrogels with Spatially Patterned Biochemical Signals to Control Cell Behavior. Adv. Mater. 2015, 27 (8), 1356-1362. DOI: 10.1002/adma.201404993.Nanofibrous.

55. Kappes, U. P.; Luo, D.; Potter, M.; Schulmeister, K.; Rünger, T. M., Short- and Long-Wave UV Light (UVB and UVA) Induce Similar Mutations in Human Skin Cells. J. Invest. Dermatol. 2006, 126 (3), 667-675. DOI: 10.1038/sj.jid.5700093.

56. Kielbassa, C.; Roza, L.; Epe, B., Wavelength dependence of oxidative DNA damage induced by UV and visible light. Carcinogenesis 1997, 18 (4), 811-816. DOI: 10.1093/carcin/18.4.811.

57. Yin, H.-L.; Wu, C.-C.; Lin, C.-H.; Chai, C.-Y.; Hou, M.-F., β1 Integrin as a Prognostic and Predictive Marker in Triple-Negative Breast Cancer. Int. J. Mol. Sci. 2016, 17 (9), 1-15. DOI: 10.3390/ijms17091432.

58. Lu, P.; Weaver, V. M.; Werb, Z., The extracellular matrix: A dynamic niche in cancer progression. J. Cell Biol. 2012, 196 (4), 395-406. DOI: 10.1083/jcb.201102147.

59. Plow, E. F.; Haas, T. A.; Zhang, L.; Loftus, J.; Smith, J. W., Ligand Binding to Integrins. J. Biol. Chem. 2000, 275 (29), 21785-21788. DOI: 10.1074/jbc.R000003200.

60. Emsley, J.; Knight, C. G.; Farndale, R. W.; Barnes, M. J.; Liddington, R. C., Structural Basis of Collagen Recognition by Integrin alpha2Beta1. Cell 2000, 101 (1), 47-56. DOI: 10.1016/S0092-8674(00)80622-4.

